# Generative frame interpolation enhances tracking of biological objects in time-lapse microscopy

**DOI:** 10.1101/2025.03.23.644838

**Authors:** Swaraj Kaondal, Arsalan Taassob, Sara Jeon, Su Hyun Lee, Henrique L. Nuñez, Bukola A. Akindipe, Hyunsook Lee, So Young Joo, Samuel M.D. Oliveira, Orlando Argüello-Miranda

**Affiliations:** Department of Plant and Microbial Biology, North Carolina State University, Raleigh, USA; Institute of Molecular Biology and Genetics, Seoul National University, Seoul, Korea; Joint School of Nanoscience and Nanoengineering, North Carolina A&T State University, Greensboro, USA

## Abstract

Object tracking in microscopy videos is crucial for understanding biological processes. While existing methods often require fine-tuning tracking algorithms to fit the image dataset, here we explored an alternative paradigm: augmenting the image time-lapse dataset to fit the tracking algorithm. To test this approach, we evaluated whether generative video frame interpolation can augment the temporal resolution of time-lapse microscopy and facilitate object tracking in multiple biological contexts. We systematically compared the capacity of Latent Diffusion Model for Video Frame Interpolation (LDMVFI), Real-time Intermediate Flow Estimation (RIFE), Compression-Driven Frame Interpolation (CDFI), and Frame Interpolation for Large Motion (FILM) to generate synthetic microscopy images derived from interpolating real images. Our testing image time series ranged from fluorescently labeled nuclei to bacteria, yeast, cancer cells, and organoids. We showed that the off-the-shelf frame interpolation algorithms produced bio-realistic image interpolation even without dataset-specific retraining, as judged by high structural image similarity and the capacity to produce segmentations that closely resemble results from real images. Using a simple tracking algorithm based on mask overlap, we confirmed that frame interpolation significantly improved tracking across several datasets without requiring extensive parameter tuning and capturing complex trajectories that were difficult to resolve in the original image time series. Taken together, our findings highlight the potential of generative frame interpolation to improve tracking in time-lapse microscopy across diverse scenarios, suggesting that a generalist tracking algorithm for microscopy could be developed by combining deep learning segmentation models with generative frame interpolation.

## Introduction

Tracking objects such as single cells and biological structures in microscopy images is a fundamental challenge of time-lapse microscopy in biological and biomedical sciences. In cell biology, tracking is required to study structures ranging from molecules to single cells that can follow complex trajectories [1,2]. Tracking biological objects remains a challenge that often requires advanced mathematical modeling for different datasets. In this work, we suggest a generalist approach to improve tracking by increasing the overlap of biological objects across frames using generative deep learning frame interpolation.

Conventional microscopy tracking pipelines begin by segmenting individual objects in each frame of a time-lapse series. Tracking then relies on linking these segmentations or “masks” over successive frames to consistently identify each object. The current standard for image segmentation is deep learning algorithms, such as U-Nets and pixel flow-directed convolutional neural networks [3, 4, 5, 6]. Once images are correctly segmented, tracking depends on the selection of an appropriate algorithm for the image dataset.

Most tracking algorithms depend on specific parameters to link the segmented masks of biological objects across frames. Tracking approaches based on the Viterbi framework rely on optimizing linking distances and assumptions about object velocity [7,8]. Advanced Bayesian or Gaussian mixture-based tracking models necessitate motion models and estimator counts [9, 10]. Other widely used tracking approaches employ global cost minimization via the Viterbi algorithm [7], probabilistic frameworks (e.g., Bayesian tracking [9]), or Kalman filtering for motion prediction [11]. These methodologies often require parameter tuning to accommodate variations in object density, overlap and occlusion, sampling frequency, and signal-to-noise ratios [1]. Mathematical approaches to overcome this complexity include the u-track methods [12] and automated tools for cell segmentation and tracking, such as ALFI [13], CellTraxx [14], ThirdPeak [15], and TrackMate [16].

Given that most tracking approaches depend on parameters such as linking distances, allowable skipped frames, or anticipated object speeds, one straightforward approach to improve tracking is to increase the sampling rate to obtain higher object overlap in consecutive frames. Experimentally, however, increasing the frequency of image acquisition can lead to phototoxicity or photobleaching, potentially perturbing biological target processes and reducing data quality [17]. A post-acquisition solution to achieve higher temporal resolution is frame interpolation, whereby information from consecutive real frames is used to produce intermediate synthetic frames in between real frames. Content-Aware Frame Interpolation (CAFI) and related algorithms use deep learning frame interpolation [17, 18, 19] for enhancing super-resolution imaging [16, 17], improve contrast in cryo-transmission electron microscopy [20, 18, 17], multi-resolution data fusion [21, 22], deconvolution [23, 24, 25], surface projection [26, 27], and denoising [28,29,30]. However, exploiting generative frame interpolation to augment microscopy time series to improve tracking remains relatively underexplored.

In this work, we explored whether augmenting image time series through generative frame interpolation simplifies the tracking of biological objects (**Fig. 1 A,B**). We evaluated whether generative deep learning-based frame interpolation methods can produce bio-realistic microscopy images to facilitate tracking across diverse microscopy datasets. We employ four deep learning frame interpolation algorithms: Latent Diffusion Model for Video Frame Interpolation (LDMVFI)[31], which leverages latent space modeling to predict and synthesize intermediate frames; Real-Time Intermediate Flow Estimation (RIFE)[32], which estimates and interpolates pixel motion between frames; Compression-Driven Frame Interpolation (CDFI)[33], which optimizes frame transitions based on data compression techniques to maintain visual coherence; and Frame Interpolation for Large Motion (FILM)[34], which addresses large-scale and rapid movements within frame sequences. Each model was tested in diverse time-lapse datasets composed of fluorescently labeled nuclei in the fungus *Ashbya gossypii*, proliferating *Escherichia coli* bacteria, proliferating yeast *Saccharomyces cerevisiae*, germinating filaments of the fungal plant pathogen *Colletotrichum acutatum*, migrating rat basophilic leukemia cancer cells, a classical video of a neutrophil chasing a bacterium, and pancreas-derived organoids.

**Figure 1.**
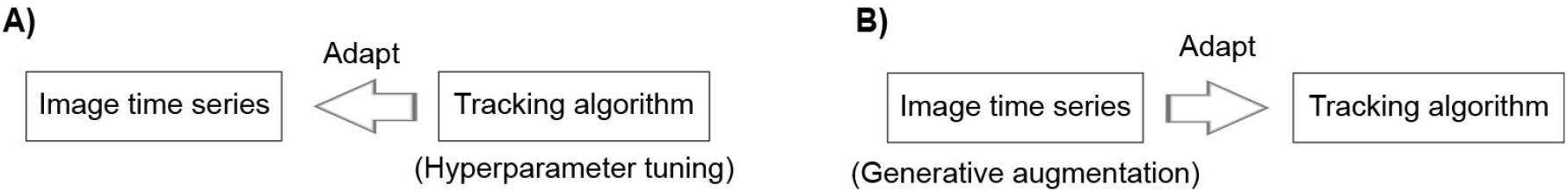
Schematic contrasting **(A)** the most common approach to tracking objects in biological time series and **(B)** the approach proposed in this work, in which the time series itself is modified using generative AI algorithms to fit the demands of a tracking algorithm without modifying the algorithm itself.

Our results show that generative frame interpolation produced bio-realistic images across all imaging modalities and biological scenarios without dataset-specific retraining. As proof of principle, we showed that the temporal augmentation of the image time series enhanced the performance of a simple mask overlap-based tracking algorithm across all tested datasets, underscoring the potential of generative frame interpolation for reaching generalist microscopy tracking.

## Results

### Deep learning interpolation generates bio-realistic images

To assess the capacity of generative frame interpolation to produce new microscopy images in between two frames from an image time series, we implemented the pre-existing off-the-shelf video frame interpolation algorithms LDMVFI [31], RIFE [32], CDFI [33], and FILM [34]. As datasets for testing interpolations, we gathered eight microscopy time series with significantly different image properties, such as single pixel entropy and the Gray-Level Co-Occurrence Statistics (GLCOS) contrast, homogeneity, and correlation (**SFig. 1 A-D, Video 1**). The time series represented intracellular structures (**Fig. 2 A**, fluorescently labeled nuclei in the fungus *Ashbya gossypii*), bacterial cells (**Fig. 2 B**, proliferating *E. coli* bacteria), budding cells (**Fig. 2 C**, proliferating yeast *Saccharomyces cerevisiae*), filamentous cells (**Fig. 2 D**, germinating spores of the fungal plant pathogen *Colletotrichum acutatum*), migrating cancer cells (**Fig. 2 E-F**, rat basophilic leukemia (RBL) cancer cells at two magnifications), immune cells (**Fig. 2 G**, a classical movie of a neutrophil chasing a bacterium) and pancreas-derived mouse organoids exposed to anticancer drugs (**Fig. 2 H**).

**Figure 2.**
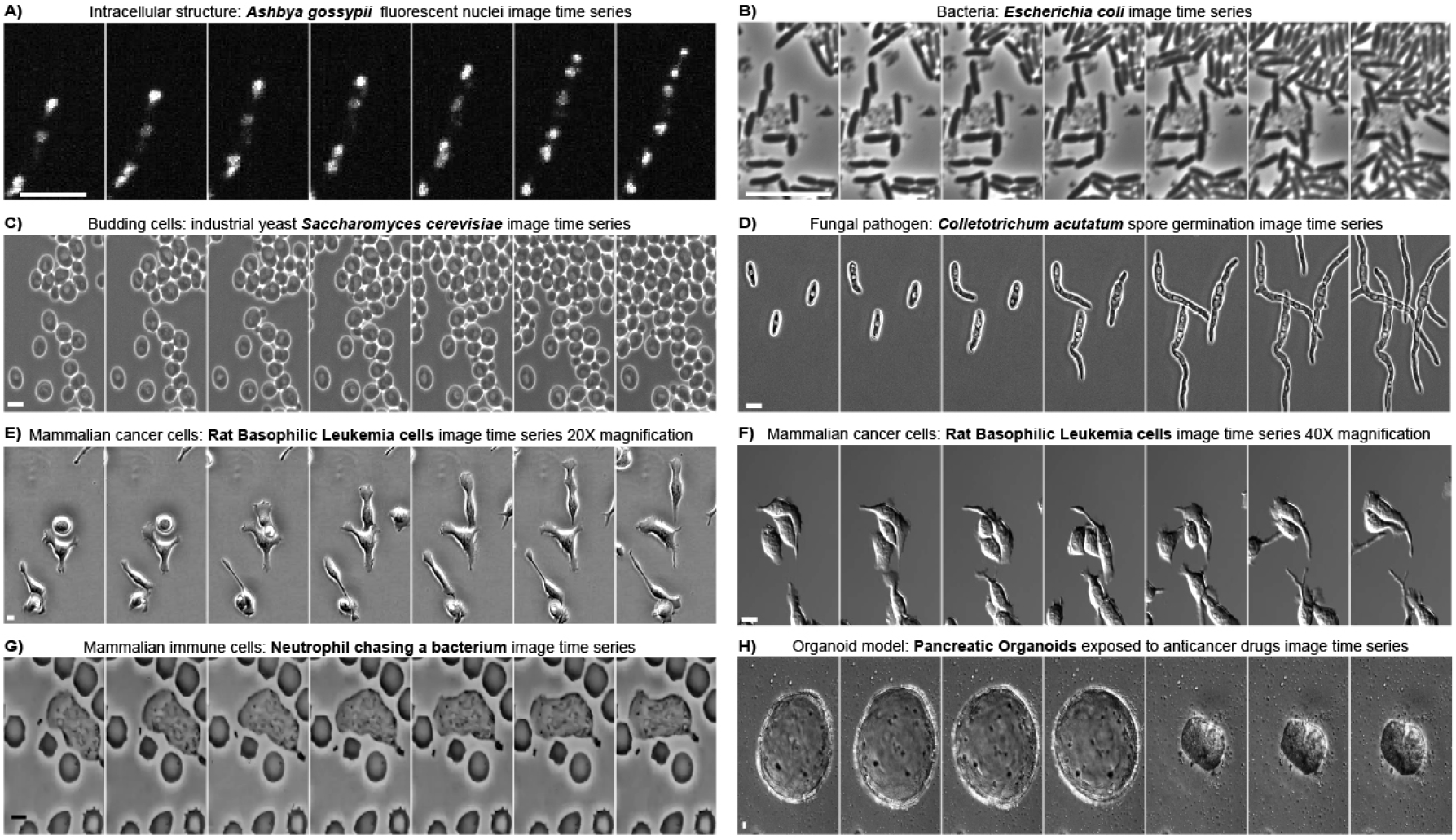
Representative image time series from time-lapse microscopy data with significantly different image properties. (**A**) intracellular structures: fluorescently labeled nuclei in the fungi *Ashbya gossypii;* (**B**) bacterial cells: proliferating *Escherichia coli* bacteria; (**C**) budding cells: proliferating yeast *Saccharomyces cerevisiae*; (**D**) filamentous cells: germinating spores of the fungal pathogen *Colletotrichum acutatum* (**E-F**) migrating cancer cells: Rat Basophilic Leukemia (RBL) cancer cells at (**E**) 20 X and (**F**) 40X magnification; (**G)** immune cells: a classical movie of a neutrophil chasing a bacterium; and (**H**) pancreas-derived organoids collapsing in response to anticancer drugs. Brightness and intensity normalized for display purposes. All scale bars = 5 µm.

To test whether frame interpolation could accurately reproduce real images, we removed the odd-numbered frames from the videos and used the even-numbered frames to produce interpolations representing the odd-numbered images. The interpolation accuracy was calculated by comparing the real odd images, as ground truth, to the interpolated odd images. The similarity between images was assessed using the structural similarity index measure (SSIM) [31] (**Fig. 3 A-H)**. We found that all models were able to produce images with great similarity across the time series as judged by SSIM values over 0.9 (**Fig. 3 I-P**). Although all SSIM values remained close to one, diffusion-based LDMFVI performed significantly lower than the rest in the fluorescent nuclei and yeast time series **(Fig 3 I,K)**. In contrast, all models showed no difference for the other image time series **(Fig. 3 J,L,M,N,O,P)**. The image’s background area percentage did not influence the interpolation quality **(SFig. 2 A)**.

**Figure 3.**
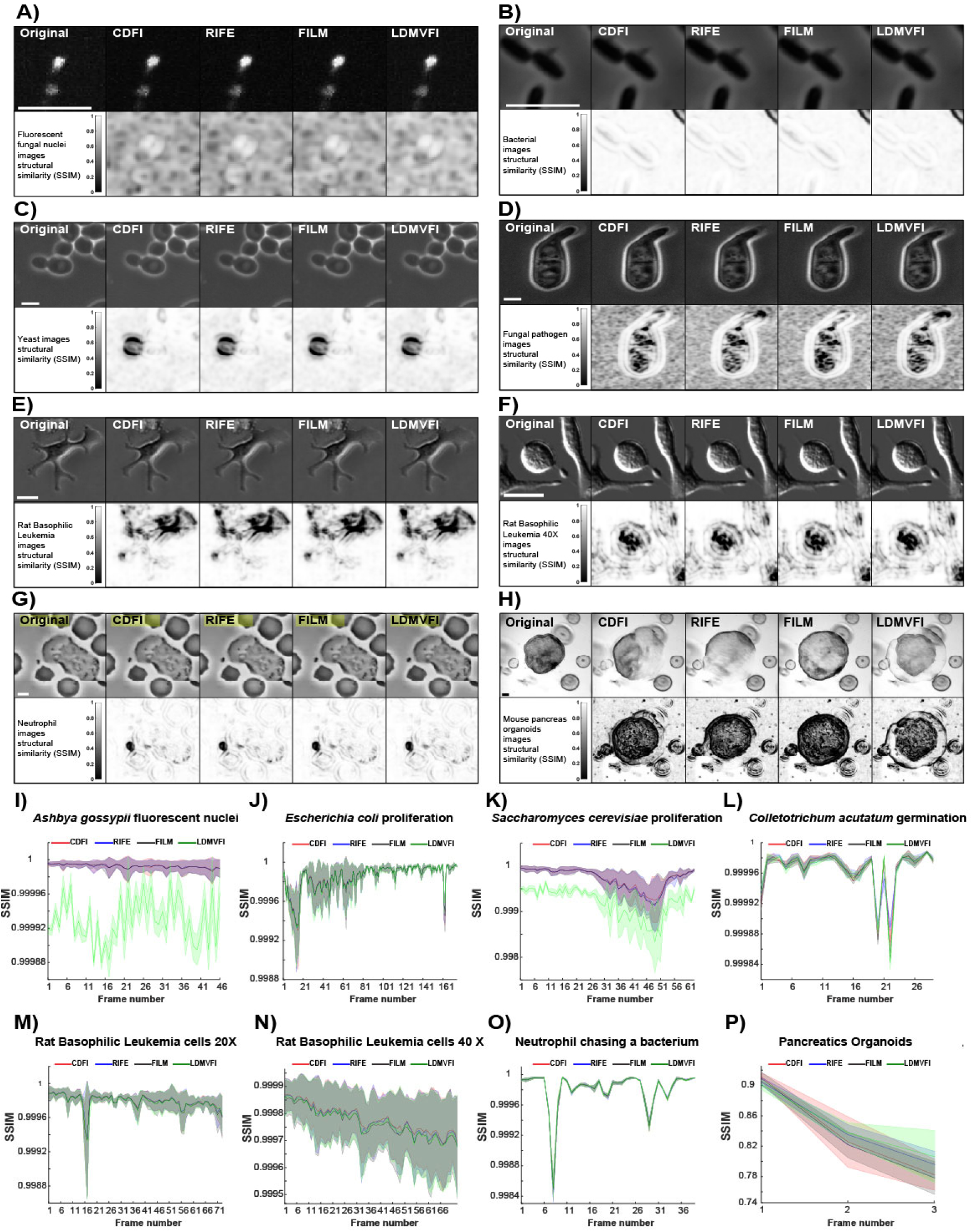
Deep learning generative frame interpolation produces images with bio-realistic structural information. **(A-H) Top:** representative images showing the interpolation results derived from using even-numbered images to reconstruct odd-numbered images with each interpolation model; **Bottom:** structural similarity heatmaps showing the areas of disagreement between the original and the interpolated image for each model as darker pixels. **(I-P)** Average SSIM values for each time point of the interpolated time series for each model and dataset compared to the original time series. Solid lines with shaded area = average plus 95% confidence intervals, n >= 4 per time point. White or dark scale bars = 5 μm.

SSIM were the highest for objects with homogenous internal features, such as fluorescent nuclei images **(Fig. 3 A)** and bacteria **(Fig. 3 B)**. The lowest SSIM regions consistently occurred when representing small extracellular objects, for instance, the growing yeast bud in **(Fig. 3 C)** or the bacterium in the neutrophil image **(Fig 3 G)**, or when representing the complex intracellular features of germinating fungal spores **(Fig 3 D)** or RBL cells **(Fig 3 E,F)**. The exception to high SSIM values was the interpolation of a low frame rate time series of pancreas organoids collapsing in response to an anti-proliferative drug **(Fig. 3 H,P)**. We concluded that deep learning frame interpolation algorithms could generate bio-realistic images across multiple biological data sets without specific retraining for microscopy.

### Synthetic interpolated images recapitulate the segmentation results of real images

Given the bio-realistic images obtained by odd-even interpolations, we tested whether interpolated images could be segmented as efficiently as real images. We assessed the segmentability of synthetic images by using the following criterion: an efficiently interpolated image should produce the same segmentation result as the original real image when using a segmentation model trained on real images.

We trained segmentation models for each data set based exclusively on real images using Cellpose [6]. We then interpolated all odd and even frames to produce time series entirely composed of interpolated images. Both the real and the fully interpolated time series were then segmented, and the resulting masks were compared using Average Precision (AP) at the single object mask level. Briefly, the AP for each object mask was calculated by dividing the total true positive (TP) pixels by the sum of the total true positive (TP) pixels plus total false positive (FP) pixels plus total false negative (FN) pixels or AP = TP / (TP + FP + FN) [6].

We found that real and interpolated images generated segmentations with very high similarity as judged by corrected average precision values above 0.7, which remained constant throughout most of the time series **(Fig. 4 A-G)**. Except for LDMFVI in the bacterial dataset **(Fig. 4 B)**, average AP values were not significantly different across models. We concluded that the segmentation of synthetic interpolated images resembled the segmentation of real images.

**Figure 4.**
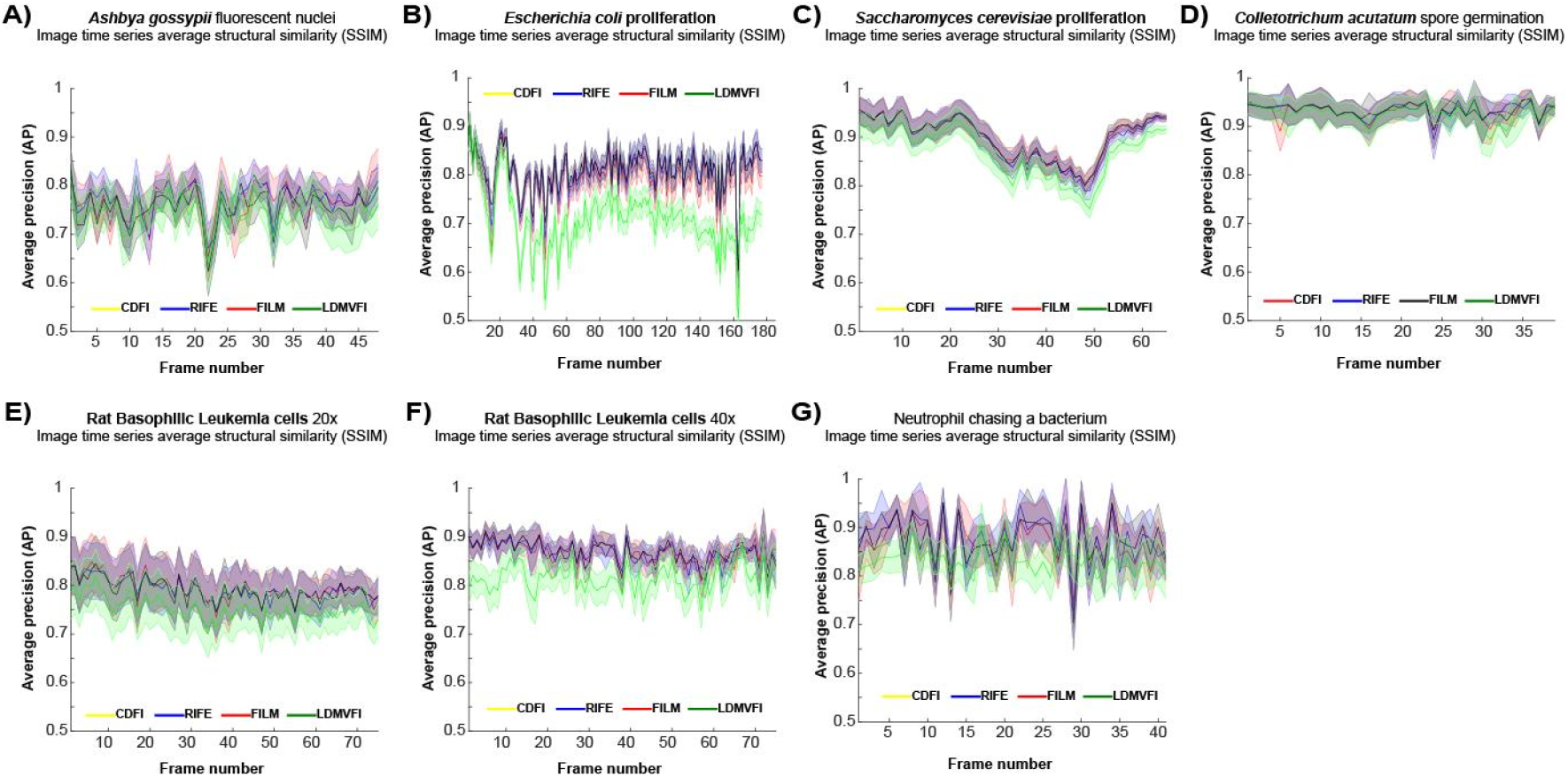
Synthetic interpolated and real images produce similar segmentation results at the single mask level. **(A-G)** Average precision values at the single mask level for each time point when comparing the segmentations from interpolated versus real images for each interpolation model. (**A**) Intracellular structures: fluorescently labeled nuclei in the fungi *A. gossypii*. (**B**) Bacterial cells: proliferating *E. coli* bacteria. (**C**) Budding cells: proliferating industrial yeast *S. cerevisiae*. (**D**) Filamentous cells: germinating spores of the fungal plant pathogen *C. acutatum*. (**E-F**) migrating cancer cells: RBL cancer cells at (**E**) 20 X and (**F**) 40X magnification. (**G)** Immune cells: a classical movie of a neutrophil chasing a bacterium. Solid lines with shaded area = average plus 95% confidence intervals, n >= 4 per time point.

### Frame interpolation recreates potential intermediate states of biological objects

The similarity in the segmentation results of real and interpolated frames suggested that interpolations between frames could generate images depicting potential intermediate states of biological objects, which could be segmented and leveraged for tracking.

To assess the capacity of the interpolation algorithms to recreate the potential intermediate states of biological objects, we generated sixteen interpolated images in between real frames across data sets **(Videos 2-9)**. Visual inspection of the 16X interpolated time series confirmed that interpolation could produce images depicting the potential intermediate positions and shapes of biological objects such as the deformation of nuclei during nuclear movements **(Fig. 5 A)**, the rotation of bacterial cells during proliferation **(Fig. 5 B)**, the growth of a budding yeast cell **(Fig. 5 C)**, and shape-shifts in the protrusions (lamellipodia) of RBL cells **(Fig. 5 D)**. At the single object level, the models showed slight variations in fine details that led to differences in the predicted masks across models, for instance, in depicting cell protrusions in RBL cells **(Fig. 5 E)**. The SSIM of the interpolated images and the AP of their segmentations decayed in comparison to the previous original image and increased in comparison to the next real image, which is consistent with interpolations producing images with intermediate segmentable information **(Fig 5 F, SFig. 3 A-F)** and structural similarity (**SFig. 3 G-L**).

**Figure 5.**
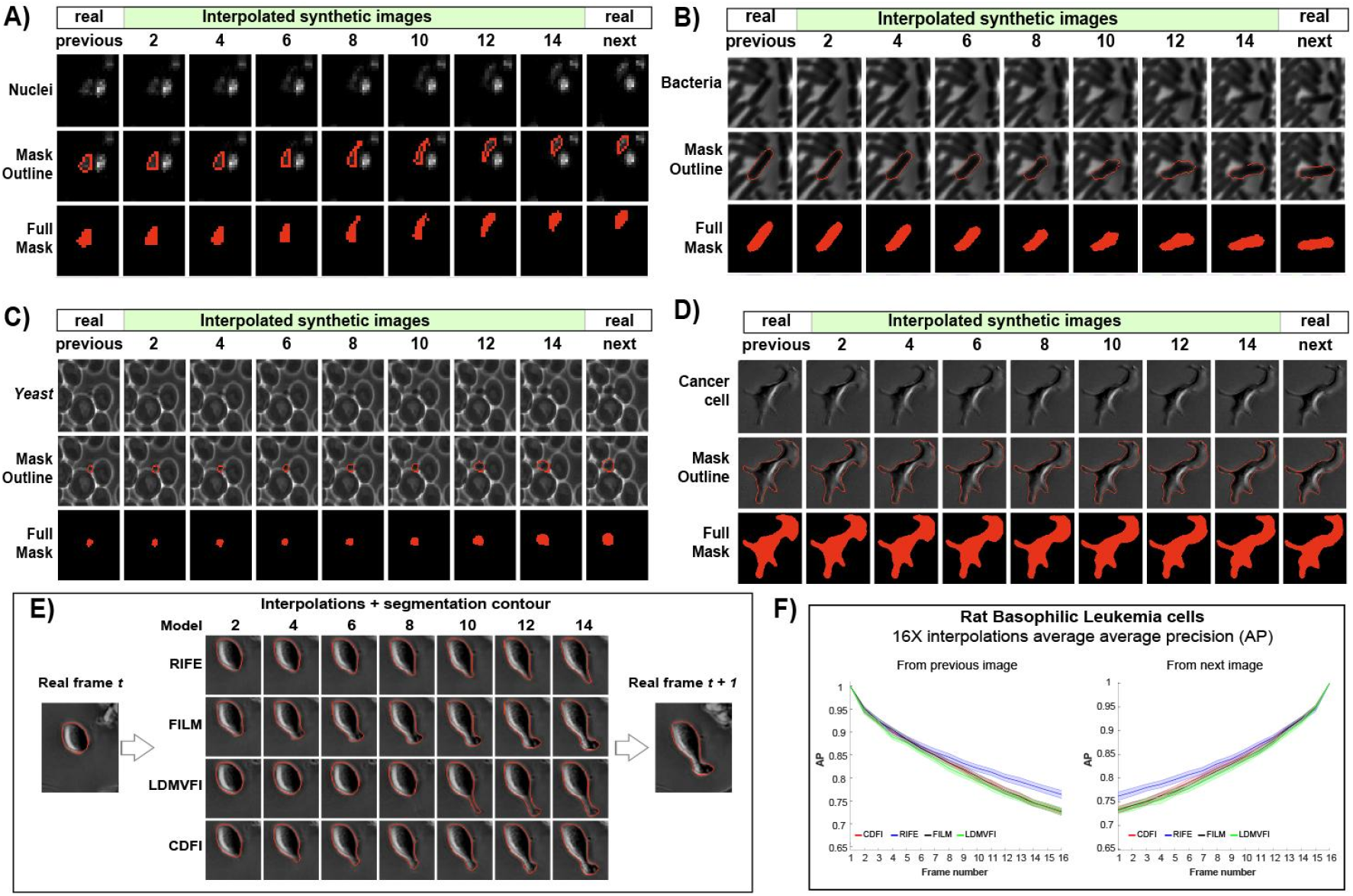
Extensive image interpolation generates potential intermediate depictions of biological objects. **(A-D)** Representative hybrid time series with real and interpolated images for **(A)** fluorescently labeled nuclei in the fungi *A. gossyipi* obtained using LDMVFI (**B**) proliferating *E. coli* bacterial cells obtained using FILM. (**C**) Budding yeast *S. cerevisiae* cells using RIFE. (**D**) shape-shifting migrating RBL cancer cells using CDFI. Top: hybrid time series with real and interpolated images. Middle: hybrid time series with mask outlines. Bottom: object’s mask across the time series. (**E**) Representative comparison of the different mask outlines obtained by interpolating two consecutive frames with different models. **(F)** Average Precision values for the segmented frames of 16 X interpolated time series for the RBL dataset. Left: AP values for each segmented interpolated frame using the segmentation of the previous real image as reference. Right: AP values for each segmented interpolated frame using the segmentation of the next real image as reference. Solid lines with shaded area = average plus 95% confidence intervals, n >= 4 per time point.

### Generating hybrid real-synthetic time series improves the tracking of biological objects

The potential of interpolated images to depict bio-realistic intermediate states of biological objects suggested that they could be used to link labeled objects across frames based solely on mask overlap. To test this hypothesis, we designed a simple tracking algorithm based on linking objects across frames by only calculating the overlap of the previous and next object mask. Using this simple approach, we tracked fluorescent nuclei, bacteria, yeast, and RBL datasets, which contained enough complex object trajectories. We segmented and tracked each dataset’s original time series and hybrid time series containing 16 interpolated images in between real images.

After tracking, the interpolated time series were down-sampled so that the number of frames matched the original time series again. We manually assessed the number of correctly assigned tracks in each dataset as a measure of tracking efficiency using a user interface that displayed each frame with a mask overlaid for each tracked object. Frame interpolation significantly increased the percentage of correctly assigned tracks **(Fig. 6 A-D, Videos 10-13)**, except for the diffusion LDMVFI model, which showed no difference with the original non-interpolated nuclei and yeast times series **(Fig. 6 A, C)**.

**Figure 6.**
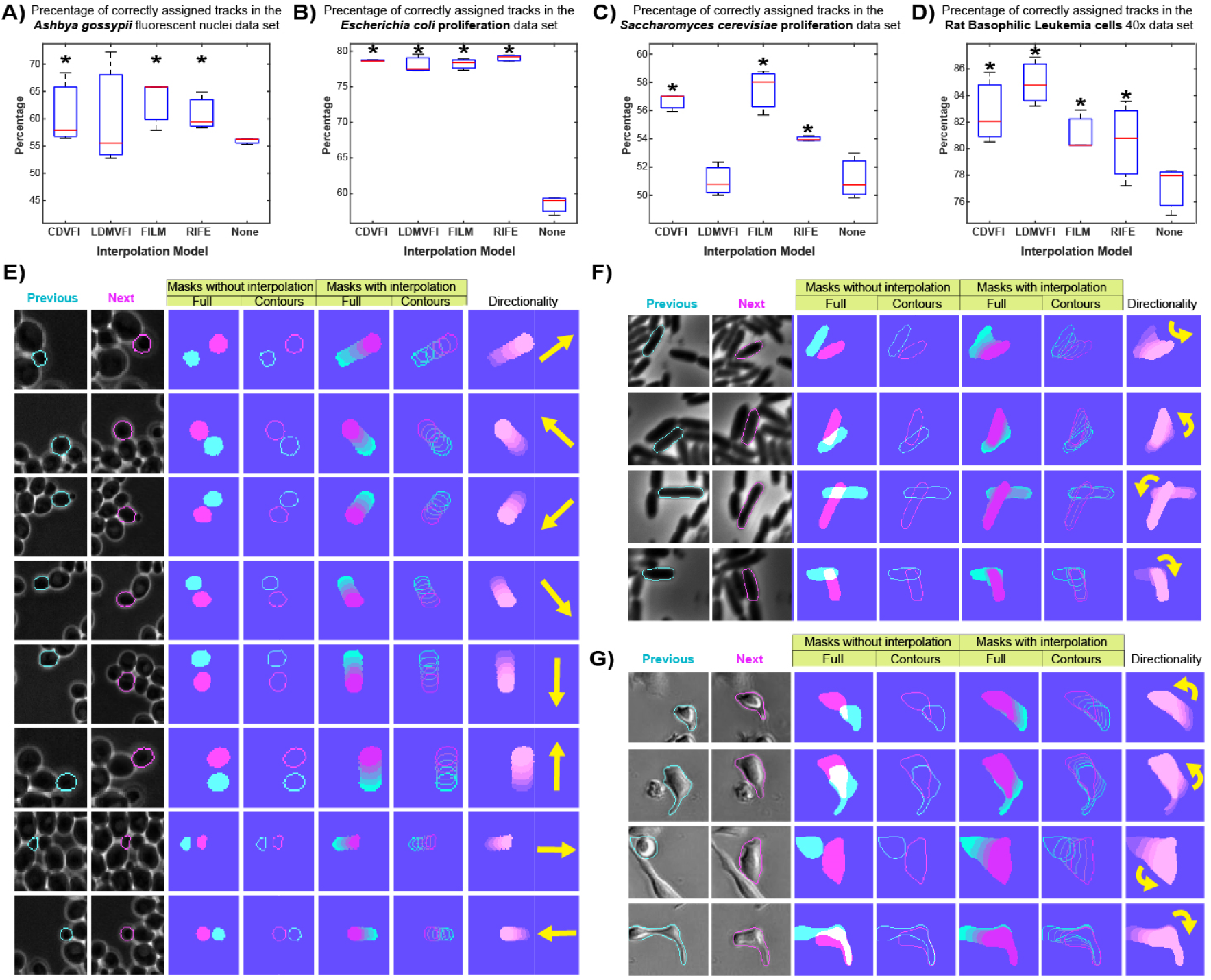
Extensive frame interpolation enhances a tracking algorithm across multiple datasets. **(A-D)** Boxplots comparison of the percentage of correctly assigned tracks with and without frame interpolation using the same tracking algorithm by mask overlap on **(A)** fluorescently labeled nuclei in the fungi *A. gossypii*; (**B**) proliferating *E. coli* bacteria; (**C**) proliferating yeast *S. cerevisiae*. (**D**) migrating rat basophilic leukemia cancer cells at 20X magnification. Box plots display data from n >= 3 biological replicates. Central mark, median; box bottom and top limit, 25th and 75th percentiles; whiskers, most extreme non-outlier values. Asterisks = p > 0.05, KS test, n >= 3. **(E)** Panel depicting examples of tracked *S. cerevisiae* cells that move in multiple directions between two consecutive frames. Columns previous and next depict the original frames with mask contours overlaid; mask overlap without and with RIFE interpolation is depicted in the columns “Full” and “Contours”; the direction of the movement is represented in the “Directionality” column. **(F)** Panel depicting examples of tracked *E. coli* cells that rotated by multiple degrees between two consecutive frames. Columns previous and next depict the original frames with mask contours overlaid; mask overlap without and with FLIM interpolation is depicted in the columns “Full” and “Contours”; the rotation direction is represented in the “Directionality” column. **(G)** Panel depicting examples of tracked Rat Basophilic Leukemia cancer cells that migrate in different directions while changing rapidly between two consecutive frames. Columns previous and next depict the original frames with mask contours overlaid; mask overlap without and with CDFI interpolation is depicted in the columns “Full” and “Contours”; the direction of migration is represented in the “Directionality” column.

Visualization of the tracks with and without interpolation confirmed that interpolation improved tracking by increasing the overlap of masks even when no overlap occurred in the original time series. This was largely independent of the direction of movement and the size of the object, as illustrated in the yeast and nuclei dataset (**Fig. 6 E, SFig. 4 A**), and regardless of rotations, as illustrated in the bacterial database **(Fig. 6 F**), and despite dramatic morphological changes as seen during cell migration in the RBL dataset (**Fig. 6 G, SFig. 4 B, C**).

## Discussion

This study demonstrates that off-the-shelf generative frame interpolation algorithms can produce bio-realistic microscopy images and improve object tracking without dataset-specific retraining. Our results indicate that generative frame interpolation algorithms trained on large video datasets learn sufficiently generic spatiotemporal priors to generate realistic intermediate frames for microscopy in diverse biological systems. This generalization echoes findings in related fields where broadly trained models, such as kernel-based interpolation techniques [35], can accurately render intermediate frames under diverse conditions without extensive fine-tuning. This suggests that, as hypothesized by Weigert et al. [17] and other frameworks [33, 23], the inclusion of generative interpolation algorithms into pre-existing microscopy tracking pipelines could be explored as an option to improve biological tracking. For instance, it is feasible to consider that frame interpolation could be used as an automated pre-processing step in workflows for CellTraxx [14], ThirdPeak [15], u-track3D [12], ALFI [13], and Trackmate [17].

The bio-realistic images obtained by interpolation, however, can not be taken as real information for the quantification of biological processes at this point. We show that although the interpolation algorithms produced images that sufficiently recreate cell contours for segmentation purposes; fine intracellular features and small moving objects in the frames differed in structural similarity across all interpolation models (**Fig. 3**). The bio-realism of such representations could be enhanced by integrating mechanistic or biological constraints into the interpolation models, as seen in biologically informed neural networks [36] and data-driven conditional deep generative networks [37].

The main disadvantages and limitations of the frame interpolation algorithms tested are (1) the need for parallel computing resources, such as graphics processor units (GPUs), (2) the decreased performance at very low sampling rates (for instance, the organoid collapse in **Fig. 3 H**), (3) variable results between diffusion-driven and the other models (**Fig. 6 A,C**). These considerations indicate that the tested interpolation algorithms could be inaccurate below a certain sampling rate limit that might be dataset-specific. In addition, the choice of interpolation model could affect the capacity of interpolation to enhance tracking. Nevertheless, our results show that under common microscopy experimental conditions and sampling rates, off-the-shelf interpolation methods produce results that can be advantageous for single-cell tracking.

Our work demonstrates that existing generative frame interpolation models have the potential to significantly enhance the tracking of biological objects in diverse time-lapse microscopy settings. Our results underscore a relatively straightforward and dataset-agnostic approach to augment microscopy time series and alleviate challenges of complex parameter tuning associated with standard tracking pipelines. The significant improvement of a simple tracking algorithm by frame interpolation across multiple datasets highlights the practical utility of integrating existing generative models into current tracking approaches. Furthermore, our results hint at the possibility of developing generalist tracking solutions for microscopy based on image time series augmentations.

## Materials and Methods

### 1. Interpolation algorithms

All interpolation algorithms were implemented as specified in their original papers using custom Python scripts to automatically direct the segmentation of times series (see 7. Code and Data availability). **Latent Diffusion Model for Video Frame Interpolation (LDMVFI) [31]:** An interpolation model that leverages latent diffusion, instead of directly processing pixel values, LDMVFI first encodes video frames into a lower-dimensional latent space, where it iteratively refines frames using a diffusion process. The model then generates intermediate frames by guiding noisy latent representations to reconstruct structures between observed frames. **Compression-Driven Frame Interpolation (CDFI)[33]:** A compact version of the Adaptive Collaboration of Flows for Video Frame Interpolation (AdaCoF) framework. It employs deformable convolutions, which adaptively reshape their kernel to align with motion patterns in image sequences. This spatially adaptive alignment facilitates fine-grained feature extraction at the pixel level, enabling the model to capture non-uniform motion. **Frame Interpolation for Large Motion (FILM)[34]:** A model designed to handle large motion in video sequences. It uses a scale-agnostic architecture that extracts multi-scale features and shares convolution weights across scales to improve the generalization of small and large motion. The model computes pixel-wise motion using a bi-directional flow estimator to capture forward and backward motion and a fusion module to combine the aligned features and generate the interpolated frames. FILM is trained with L1 loss, perceptual loss, and style loss to enhance image sharpness, texture fidelity, and perceptual quality. **Real-Time Intermediate Flow Estimation (RIFE)[32]:** A model developed for rapid frame interpolation by estimating optical flow to capture motion between frames.

### 2. Image time series data sets

#### 2a. Bacterial time series

*Escherichia coli* cells (AddGene) were cultured as a monolayer in PDMS microfluidic chip infused in media using a syringe pump. Media was infused through the chip at 1.5 μL/min for 24 hours as the chip was imaged in an inverted microscope (Nikon Eclipse Ti2) equipped with a 100x Apo (1.49 NA, oil) objective, external phase contrast system with an ECCD camera (DR-328G-CO2-SIL Clara, Andor). NIS-Elements software (Nikon Version 5.30.00) was used to automate image acquisition by taking images under phase contrast every 5 minutes. Cells were kept at 37 °C inside a cube incubation system (Incubator CH.HC5.SAT Full Enclosure, In Vivo Scientific, USA) and allowed to grow at a constant growth rate during image acquisition for 18-155h (i.e., 1-6.5 days).

#### 2b. Fungal time series

Experiments were carried out with *Saccharomyces cerevisiae* strain W303 or *Colletotrichum acutatum* (kindly provided by Dr. Tika Adhikari, NCSU, USA) grown under microfluidic conditions as described in [38]. Microfluidics experiments were performed on an automated Zeiss Observer Z1 microscope controlled by ZEN pro software and with temperature control (Zeiss), and a CellASIC ONIX microfluidics system maintained at 25 °C. Images were acquired using a 40X Zeiss EC Plan-Neofluar 40X 1.3 NA oil Ph 3 M27 immersion objective. *Ashbya gossypii* time series were kindly provided by the Gladfelter Laboratory (Duke, USA); imaging and cell culturing conditions are described in [40].

#### 2c. Mammalian single-cell datasets

The Neutrophil chasing a bacterium movie was downloaded from: https://embryology.med.unsw.edu.au/embryology/index.php/Movie_-_Neutrophil_chasing_bacteria. The Original video was a 16-mm movie by the late David Rogers at Vanderbilt University (circa 1950’s)[39]. The movie depicts a human neutrophil following a *Staphylococcus aureus* bacterium by chemotaxis through red blood cells on a blood film slide. The RBL movies were kindly provided by the Wu Laboratory (Yale, USA); imaging and cell culturing conditions are described in [41].

#### 2d. Mouse-Derived and Patient-Derived Pancreatic Organoid time series

Mouse-derived pancreatic organoids were established from lysed pancreatic ductal cells and cultured in Matrigel matrix (Corning) using an appropriate growth medium modified from [42]. All animal materials were prepared and handled in South Korea and approved by the Institutional Animal Care and Use Committee (IACUC) of Seoul National University (Approval No. **SNU-191007-4-2**). Patient-derived pancreatic organoids were obtained from endoscopic ultrasound-guided fine-needle biopsy (FNB) samples. Tissue samples were provided by Seoul Samsung Medical Center in accordance with Institutional Review Board (IRB) guidelines (Approval No. **E2106/001-003**). The samples were embedded in Matrigel (Corning) matrix and maintained in an appropriate culture medium modified from [43]. Both mouse-derived and patient-derived pancreatic organoids were treated with either DMSO or an anticancer drug after organoid formation. Live-cell imaging of patient-derived organoids was done after treatment for 42 hours at 1-hour intervals, while mouse-derived organoids were imaged for 36 hours at 6-hour intervals. Imaging was conducted using the THUNDER Imager 3D Live Cell and 3D Cell Culture microscope (Leica Microsystems).

#### 3. Generating dataset-specific cellpose segmentation models

Dataset-specific cellpose segmentation models were obtained through iterative training on manually labeled images from each data set. Training and retraining cycles stop when the models achieve AP > 0.9 on the test dataset following [6, 44]. A MATLAB script was used to evaluate AP for each mask based on the correspondence at the single pixel between a reference mask derived from a real image and the tested masks derived from interpolated images (see 7. Code and Data availability). All segmentations, interpolations, and training were performed on a Dual Intel Xeon Silver 4216 (2.1GHz,3.2GHz Turbo, 16C,9.6GT/s 2UPI, 22MB Cache, HT (100W) equipped with an Nvidia Quadro RTX5000, 16GB, 4DP, VirtualLink (XX20T) outfitted with 64GB 4×16GB DDR4 2933MHz RDIMM ECC Memory and a 2TB primary NVMe SSD.

#### 4. Image pre-processing for interpolation and segmentation

All frames were converted to three-layered (R, G, B) 8-bit depth PNGs before interpolation according to the specifications of each interpolation algorithm. For Fig3, a custom Python script enabled the generation of interpolated frames by taking time series based on odd or even frames only, generating synthetic odd or even frames. A modification of this script enables the generation of sixteen interpolated frames in between two real frames, as used for figures 4-6. The script takes as input the input and output directories and the number of frames to interpolate (see 7. Code and Data availability). In addition, the FLIM model also requires the path to a pre-trained model as an additional input to the interpolation script. To segment images using cellpose, real and interpolated images were turned into single-layered 16-bit depth uncompressed Tiff files. Segmented images were also processed in the same format for calculating AP values.

#### 5. A simple mask-overlap-based tracking algorithm

The simple Python tracking algorithm based on mask overlap was based on the following principles. After cellpose segmentation produces masks with individual indexes in each image, our algorithm projects each mask from a starting frame onto the next segmented image and identifies the cell mask with the highest overlap. Once identified, the index of the cell mask in the next image is replaced by the index of the cell mask in the previous image, forcing each cell/object to acquire a unique index throughout the time series. The algorithm has only one hyperparameter to be adjusted, the variable “disk_size” which is used to define the size of the objects to be filtered as small segmentation artifacts; disk_size = 3 if the cell area is close to 500 pixels, and disk_size = 6 if the average cell area is close to 2000 pixels (see 7. Code and Data availability).

## 6. Statistical analyses

Statistical analyses were performed on biological replicates within each time series. Differences between populations of biological replicates were evaluated using the Kolmogorov-Smirnov tests *kstest2()*, with significance set at p < 0.05. Line plots to test time series correspond to the average of biological replicates surrounded by the 95 % confidence intervals represented as shaded area. Box plots represented the median as the central mark, the 25^th^ and 75^th^ percentiles as the bottom and top limits of the box, and the most extreme non-outlier values as whiskers.

## 7. Code and Data availability

All original Python and MATLAB code employed in image analysis, interpolation, segmentation, tracking, and figure display is freely available: https://github.com/MirandaLab/GEN_interpolation_microscopy for LDMVFI diffusion model implementation, and https://github.com/MirandaLab/GEN_AI_Interpolation_Microscopy for other models. Specific instructions to install each model from its source are described, as well as our scripts to trigger interpolation of complete and down-sampled time series, *python interpolate_series*.*py* and *interpolate_between_series*.*py*, respectively. The AP evaluation MATLAB script and the Python mask-overlap algorithm are in https://github.com/MirandaLab/GEN_AI_Interpolation_Microscopy/tree/main/Masks_AP_Evaluation and https://github.com/MirandaLab/GEN_AI_Interpolation_Microscopy/tree/main/Simple_Mask_Overlap_tracking_algorithm, respectively. Toy Dataset “Pos13_1_B” to try the tracking algorithm can be downloaded from the Dryad repository: https://doi.org/10.5061/dryad.3bk3j9kw0.

## Acknowledgments

We thank Shengping Xiao and Min Wu (Yale, USA) for providing the RBL time series, and Grace McLaughlin and Amy Gladfelter (Duke, USA) for providing the *Ashbya gossypii* time series. This work was supported by grant R00GM135487 from the National Institute of General Medical Sciences of the National Institutes of Health, USA; the RISF award of the Research Innovation Science Funds from ORI/KIETS NCSU, the external grant award of the National Institute for Theory and Mathematics in Biology (NITMB); and the Research Capacity Fund (HATCH), project award no. NC02877-7002454 from the U.S. Department of Agriculture’s National Institute of Food and Agriculture to O.A.M. Organoid works were supported by AI-Bio Research Grant through Seoul National University (0413-20230047) to H.L. The authors declare no competing financial interests. In memoriam Andreas Doncic.

## References

[1] Cell Tracking Challenge; n.d. Accessed January 16, 2025. Available from: https://celltrackingchallenge.net/.

[2] Maška M, Ulman V, Svoboda D, Matula P, Matula P, Ederra C, et al. A benchmark for comparison of cell tracking algorithms. Bioinformatics. 2014;30(11):1609–17.

[3] Lugagne JB, Lin H, Dunlop MJ. DeLTA: Automated cell segmentation, tracking, and lineage reconstruction using deep learning. PLoS Computational Biology. 2020;16(7):e1007673.

[4] O’Connor OM, Alnahhas RN, Lugagne JB, Dunlop MJ. DeLTA 2.0: A deep learning pipeline for quantifying single-cell spatial and temporal dynamics. PLoS Computational Biology. 2022;18(2):e1009797.

[5] Cutler KJ, Stringer C, Lo TW, Rappez L, Stroustrup N, Brook Peterson S, et al. Omnipose: a high-precision morphology-independent solution for bacterial cell segmentation. Nature methods. 2022;19(11):1438–48.

[6] Stringer C, Wang T, Michaelos M, Pachitariu M. Cellpose: a generalist algorithm for cellular segmentation. Nature methods. 2021;18(1):100–6.

[7] Magnusson KEG, Jaldén J, Gilbert PM, Blau HM. Global linking of cell tracks using the Viterbi algorithm. IEEE Transactions on Medical Imaging. 2014;34(4):911–9.

[8] Tian C, Yang C, Spencer SL. EllipTrack: A global-local cell-tracking pipeline for 2D fluorescence time-lapse microscopy. Cell Reports. 2020;32(7).

[9] Ulicna K, Vallardi G, Charras G, Lowe AR. Automated deep lineage tree analysis using a Bayesian single-cell tracking approach. Frontiers in Computational Science. 2021;3.

[10] Chou TCea. Instant processing of large-scale image data with FACT, a real-time cell segmentation and tracking algorithm. Cell Reports Methods. 2023;3(1):100636.

[11] Tinevez JYea. TrackMate: An open and extensible platform for single-particle tracking. Nature Methods. 2017;115:80–90.

[12] Roudot P, Legant WR, Zou Q, Dean KM, Isogai T, Welf ES, et al. u-track3D: Measuring, navigating, and validating dense particle trajectories in three dimensions. Cell Reports Methods. 2023;3(12).

[13] Antonelli J, Kumar R, Sharma D, et al. ALFI: A label-free dataset for differential interference contrast microscopy tracking. Nature Communications. 2023;14:5678.

[14] Holmes J, Harrison A, Nguyen C, et al. CellTraxx: An automated tool for high-throughput migration assays in phase-contrast images. Scientific Reports. 2023;13:12345.

[15] Müller T, Meiser E, Engstler M. ThirdPeak: A flexible tool designed for the robust analysis of two- and three-dimensional tracking data. Communications Biology. 2024;7:1683.

[16] Ershov D, Phan MS, Pylvänäinen JW, Rigaud SU, Blanc LL, Charles-Orszag A, et al. TrackMate 7: Integrating state-of-the-art segmentation algorithms into tracking pipelines. Nature Methods. 2022;19:829–32.

[17] Weigert M, et al. Content-aware image restoration: pushing the limits of fluorescence microscopy. Nature Methods. 2018;15:1090–7.

[18] Ulman V, Maška M, Magnusson KEG, Ronneberger O, Haubold C, et al. An objective comparison of cell-tracking algorithms. Nature Methods. 2017;14:1141–52.

[19] Maška M, Ulman V, Delgado-Rodriguez P, de Mariscal EG, Nečasová T, et al. The Cell Tracking Challenge: 10 years of objective benchmarking. Nature Methods. 2023;20:1010–20.

[20] Jiang J, Khan A, Shailja S, Belteton SA, et al. Segmentation, tracking, and subcellular feature extraction in 3D time-lapse images. Scientific Reports. 2023;13:3483.

[21] Gustafsson MG. Surpassing the lateral resolution limit by a factor of two using structured illumination microscopy. Journal of microscopy. 2000;198(2):82–7.

[22] Müller M, et al. Open-source image reconstruction of super-resolution structured illumination microscopy data in ImageJ. Nature Communications. 2016;7:10980.

[23] Richardson WH. Bayesian-based iterative method of image restoration. J Opt Soc Am. 1972;62:55–69.

[24] Arigovindan M, et al. High-resolution restoration of 3D structures from widefield images with extreme low signal-to-noise-ratio. Proceedings of the National Academy of Sciences. 2013;110:17344–9.

[25] Preibisch S, et al. Efficient Bayesian-based multiview deconvolution. Nat Methods. 2014;11:645–8.

[26] Blasse C, et al. PreMosa: extracting 2D surfaces from 3D microscopy mosaics. Bioinformatics. 2017;33:2563–9.

[27] Shihavuddin A, et al. Smooth 2D manifold extraction from 3D image stack. Nature Communications. 2017;8:15554.

[28] Buades A, et al. A non-local algorithm for image denoising. Proceedings of IEEE Conference on Computer Vision and Pattern Recognition. 2005:60–5.

[29] Dabov K, et al. Image denoising by sparse 3-D transform-domain collaborative filtering. IEEE Transactions on Image Processing. 2007;16:2080–95.

[30] Morales-Navarrete H, et al. A versatile pipeline for the multi-scale digital recon-struction and quantitative analysis of 3D tissue architecture. eLife. 2015;4:e11214.

[31] Danier D, Zhang F, Bull D. Ldmvfi: Video frame interpolation with latent diffusion models. In: Proceedings of the AAAI Conference on Artificial Intelligence. vol. 38; 2024. p. 1472–80.

[32] Huang Z, Zhang T, Heng W, Shi B, Zhou S. Real-time intermediate flow estimation for video frame interpolation. In: European Conference on Computer Vision. Springer; 2022. p. 624–42.

[33] Ding T, Liang L, Zhu Z, Zharkov I. Cdfi: Compression-driven network design for frame interpolation. In: Proceedings of the IEEE/CVF conference on computer vision and pattern recognition; 2021. p. 8001–11.

[34] Reda F, Kontkanen J, Tabellion E, Sun D, Pantofaru C, Curless B. Film: Frame interpolation for large motion. In: European Conference on Computer Vision. Springer; 2022. p. 250–66.

[35] Briedis KM, Djelouah A, Ortiz R, Meyer M, Gross M, Schroers C. Kernel-Based Frame Interpolation for Spatio-Temporally Adaptive Rendering. In: ACM SIG-GRAPH 2023 Conference Proceedings; 2023. p. 1–11.

[36] Lagergren JH, Nardini JT, Baker RE, Simpson MJ, Flores KB. Biologically-informed neural networks guide mechanistic modeling from sparse experimental data. PLoS computational biology. 2020;16(12):e1008462.

[37] Yuan H, Cai L, Wang Z, Hu X, Zhang S, Ji S. Computational modeling of cellular structures using conditional deep generative networks. Bioinformatics. 2019;35(12):2141–9.

[38] Kociemba J, Jørgensen ACS, Tadić N, Harris A, Sideri T, Chan WY, et al. Multi-signal regulation of the GSK-3β homolog Rim11 controls meiosis entry in budding yeast. The EMBO Journal. 2024:1–31.

[39] Anderson, Cori A., Eser, Umut, Korndorf, Therese, Borsuk, Mark E., Skotheim, Jan M., Gladfelter, Amy S. Nuclear Repulsion Enables Division Autonomy in a Single Cytoplasm. Current Biology. 23,20:1999-2010

[40] Embryology U. Movie - Neutrophil chasing bacteria; n.d. Accessed: January 23, 2025. Available from: https://embryology.med.unsw.edu.au/embryology/index.php/Movie_-_Neutrophil_chasing_bacteria.

[41] Xiao S, Tong C, Yang Y, Wu M. Mitotic cortical waves predict future division sites by encoding positional and size information. Developmental Cell. 2017;43(4):493–506.

[42] Huch M, Bonfanti P, Boj SF, Sato T, Loomans CJ, Van De Wetering M, et al. Unlimited in vitro expansion of adult bi-potent pancreas progenitors through the Lgr5/R-spondin axis. The EMBO journal. 2013;32(20):2708–21.

[43] Boj SF, Hwang CI, Baker LA, Chio IIC, Engle DD, Corbo V, et al. Organoid models of human and mouse ductal pancreatic cancer. Cell. 2015;160(1):324–38.

[44] Lee BD, Gitter A, Greene CS, Raschka S, Maguire F, Titus AJ, et al. Ten quick tips for deep learning in biology. PLoS computational biology. 2022;18(3):e1009803.

