## Supplemental Figures for "Generative frame interpolation enhances tracking of biological objects in time-lapse microscopy"

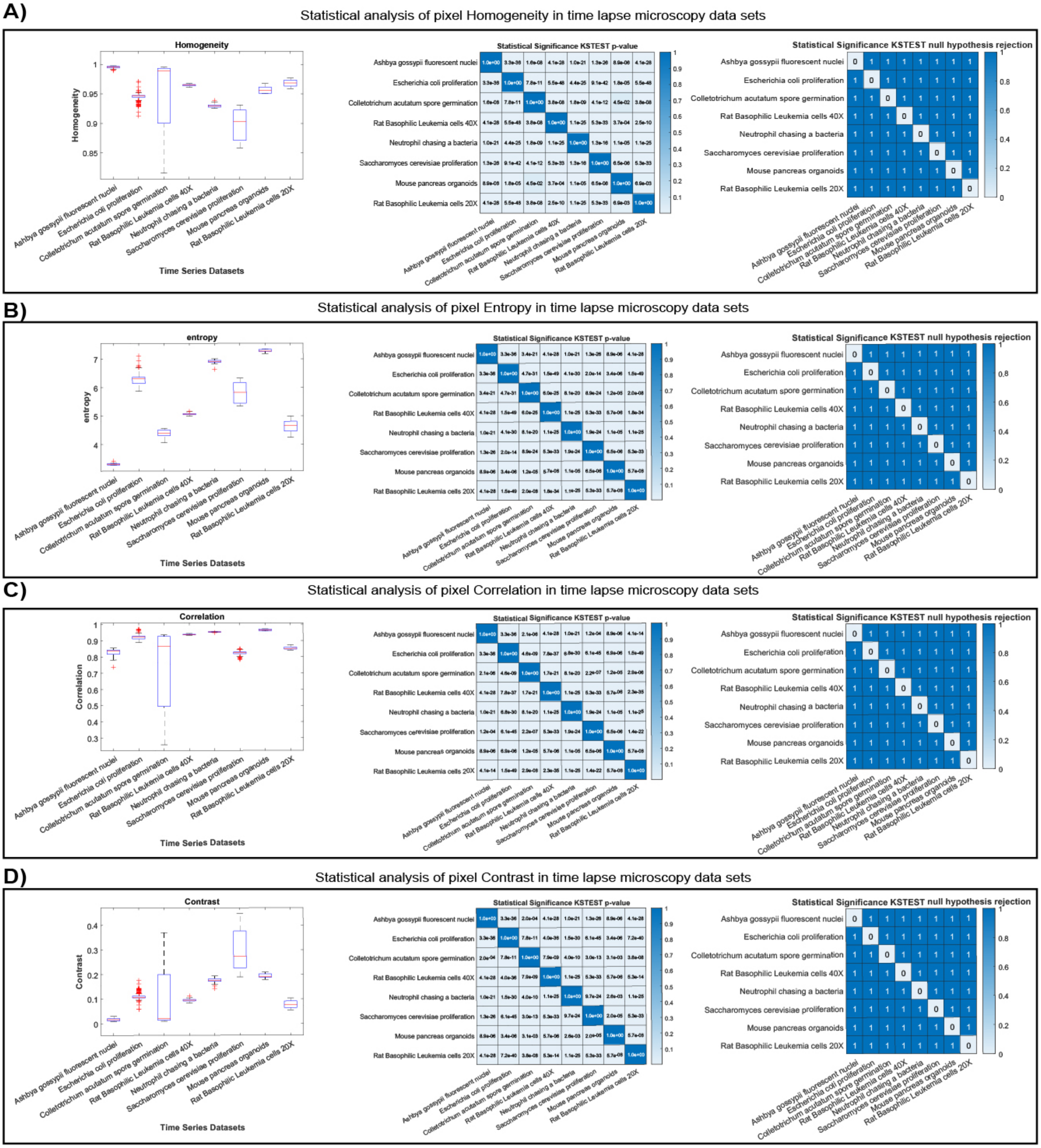


**Supplemental Figure 1. Quantification of differences in image properties across datasets displayed in figure 1** for pixel **(A)** homogeneity **(B)** entropy **(C)** correlation and **(D)** contrast. Left: Boxplot comparison of the corresponding average image properties across time-lapse data datasets. Center: Heatmaps displaying the statistical significance of the pairwise comparisons of average image properties for each dataset. Right: null hypothesis rejection heatmap. Box plots display data from biological replicates: central mark, median; box bottom and top limit, 25th and 75th percentiles; whiskers, most extreme non-outlier values. Asterisks = p > 0.05, KS test with Null hypothesis = no difference in the measured variable; n > 3.

A)

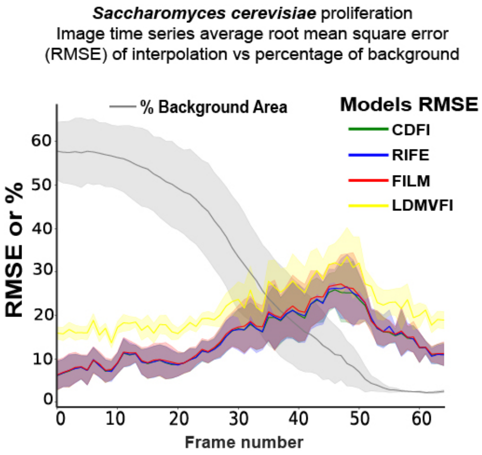


**Supplemental Figure 2. Deep learning generative frame interpolation produces images with bio-realistic structural information. (A)** Colored time series: root mean square of each interpolated image using the corresponding real image as reference. Gray time series: percentage of background in the corresponding images. The time series shows *Saccharomyces cerevisiae* cells that grow to saturation in the field of view over time (See supplemental Video 1). Solid lines with shaded area = average plus 95% confidence intervals, n >= 4 per time point.

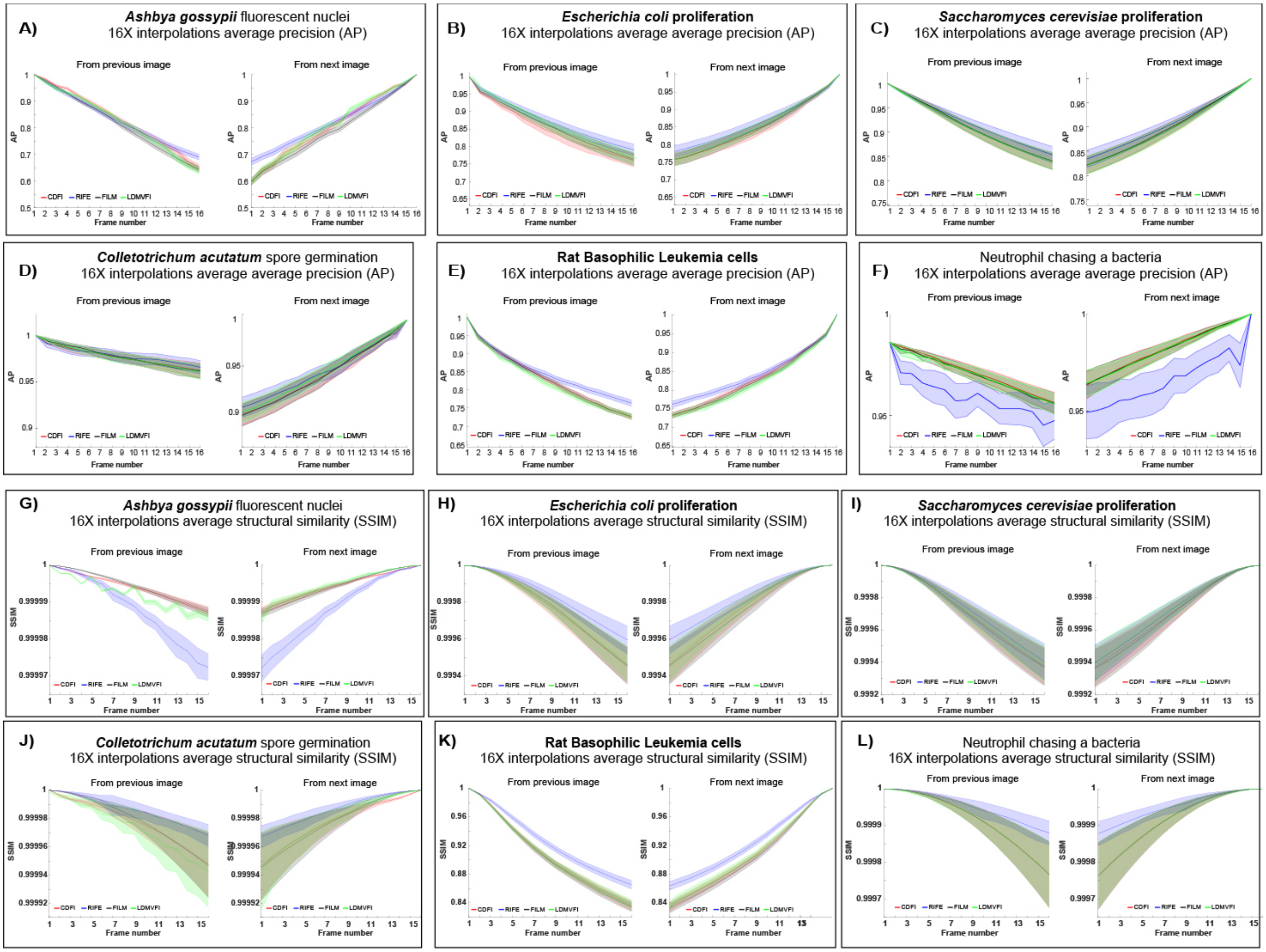


**Supplemental Figure 3**. **Extensive image interpolation generates potential intermediate depictions of biological objects**. **(A-F)** Average AP values for each frame of the 16X interpolated time series for each model and dataset. Left: AP calculations using the previous real image as reference. Right: AP values using the next real image as reference. **(G-L)** Average Precision values for the segmented frames of 16 X interpolated time series for each model and dataset. Left: SSIM values for each segmented interpolated frame using the segmentation of the previous real image as reference. Right: SSIM values for each segmented interpolated frame using the segmentation of the next real image as reference. Solid lines with shaded area = average plus 95% confidence intervals, n >= 4 per time point.

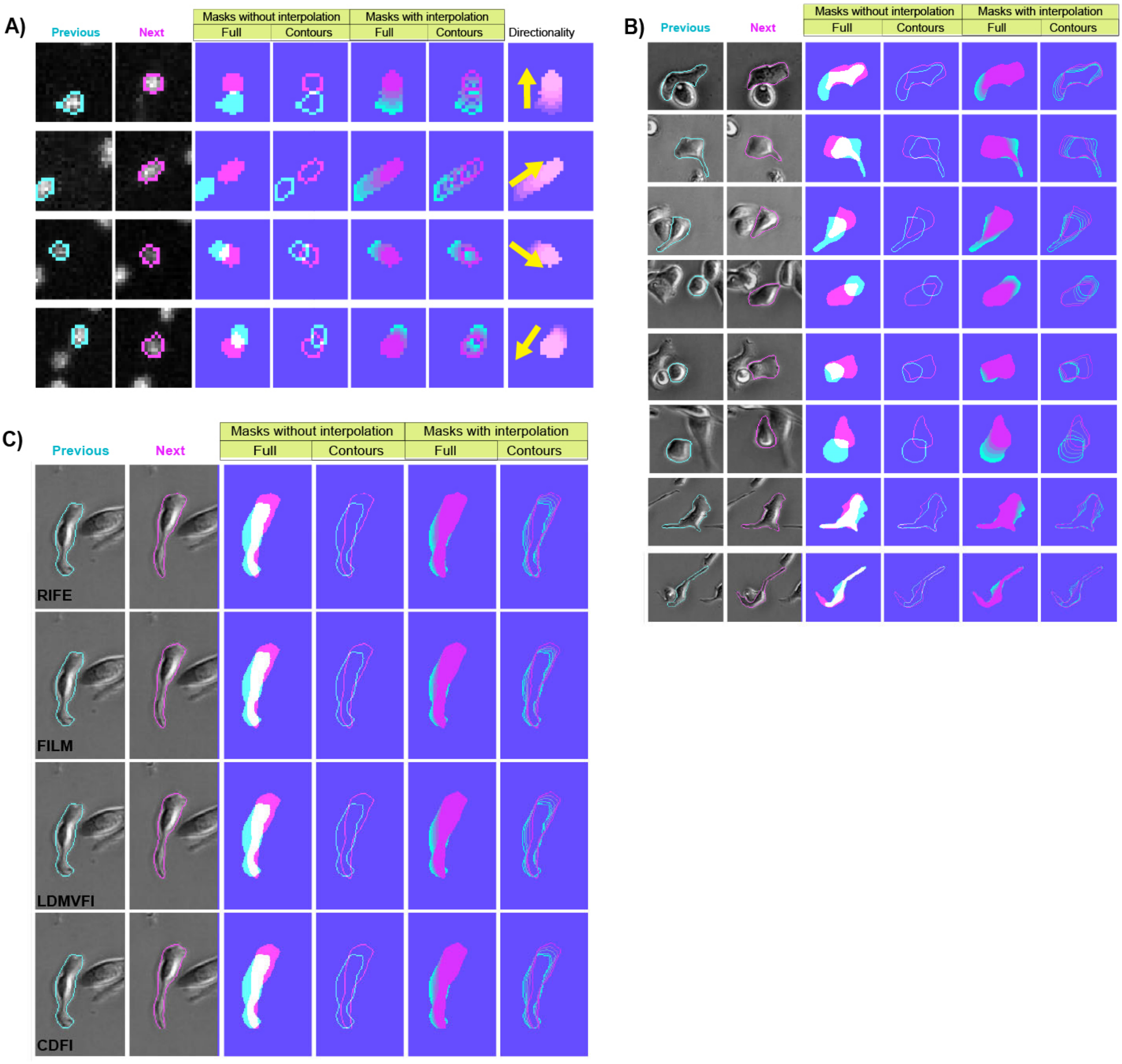


**Supplemental Figure 4. Extensive frame interpolation recreates potential intermediate states of biological objects.** **(A)** Panel depicting examples of tracked *Ashbya gossipy* fluorescently labeled nuclei that move in multiple directions between two consecutive frames. Columns previous and next depict the original frames with mask contours overlaid; mask overlap without and with LDMVFI interpolation is depicted in the columns "Full" and "Contours"; the direction of the movement is represented in the "Directionality" column. **(B)** Panel depicting examples of tracked Rat Basophilic Leukemia cancer cells that migrate in shifted shape between two consecutive frames. Columns previous and next depict the original frames with mask contours overlaid; mask overlap without and with RIFE interpolation is depicted in the columns "Full" and "Contours". **(C)** Representative Comparison of the segmentation Rat Basophilic Leukemia cancer cells images obtained through different interpolation algorithms. Columns previous and next depict the original interpolated frames with mask contours overlaid; mask overlap without and with interpolation is depicted in the columns "Full" and "Contours".

### Supplemental Video legends

**Video 1. Representative time-lapse microscopy time series. Top from left to right:** fluorescently labeled nuclei in the fungus *Ashbya gossypii;* proliferating *Escherichia coli* bacteria; proliferating yeast *Saccharomyces cerevisiae*; and germinating spores of the fungal pathogen *Colletotrichum acutatum*. **Bottom from left to right:** Rat Basophilic Leukemia (RBL) cancer cells at 20 X magnification; RBL cancer cells at 40 X magnification; a classical movie of a neutrophil chasing a bacteria; and pancreas-derived organoids collapsing in response to anticancer drugs. Brightness normalized for collective display purposes. All scale bars are displayed in Figure 2.

**Video 2. Representative time-lapse microscopy time series of fluorescently labeled nuclei in the fungi *Ashbya gossypii.* Top right:** Real time series played one frame at a time. **Top right:** Hybrid time series in which 16 FILM-interpolated images were generated between real frames. **Bottom left:** Hybrid time series in which 16 CDFI-interpolated images were generated between real frames. **Bottom right:** Hybrid time series in which 16 LDMVFI -interpolated images were generated between real frames. Brightness normalized across the time series for display purposes.

**Video 3. Representative time-lapse microscopy time series of proliferating *Escherichia coli* bacteria**. **Top right:** Real time series played one frame at a time. **Top right:** Hybrid time series in which 16 FILM-interpolated images were generated between real frames. **Bottom left:** Hybrid time series in which 16 CDFI-interpolated images were generated between real frames. **Bottom right:** Hybrid time series in which 16 LDMVFI -interpolated images were generated between real frames. Brightness normalized across the time series for display purposes.

**Video 4. Representative time-lapse microscopy time series proliferating yeast *Saccharomyces cerevisiae.* Top right:** Real time series played one frame at a time. **Top right:** Hybrid time series in which 16 FILM-interpolated images were generated between real frames. **Bottom left:** Hybrid time series in which 16 CDFI-interpolated images were generated between real frames. **Bottom right:** Hybrid time series in which 16 LDMVFI -interpolated images were generated between real frames. Brightness normalized across the time series for display purposes.

**Video 5. Representative time-lapse microscopy time series of germinating spores of the fungal pathogen *Colletotrichum acutatum*. Top right:** Hybrid time series in which 16 FILM-interpolated images were generated between real frames. **Bottom left:** Hybrid time series in which 16 CDFI-interpolated images were generated between real frames. **Bottom right:** Hybrid time series in which 16 LDMVFI -interpolated images were generated between real frames. Brightness normalized across the time series for display purposes.

**Video 6. Representative time-lapse microscopy time series of Rat Basophilic Leukemia (RBL) cancer cells at 40 X magnification. Top right:** Hybrid time series in which 16 FILM-interpolated images were generated between real frames. **Bottom left:** Hybrid time series in which 16 CDFI-interpolated images were generated between real frames. **Bottom right:** Hybrid time series in which 16 LDMVFI -interpolated images were generated between real frames. Brightness normalized across the time series for display purposes.

**Video 7. Representative time-lapse microscopy time series of Rat Basophilic Leukemia (RBL) cancer cells at 20 X magnification. Top right:** Hybrid time series in which 16 FILM-interpolated images were generated between real frames. **Bottom left:** Hybrid time series in which 16 CDFI-interpolated images were generated between real frames. **Bottom right:** Hybrid time series in which 16 LDMVFI -interpolated images were generated between real frames. Brightness normalized across the time series for display purposes.

**Video 8. Representative time-lapse microscopy time series of a classical movie of a neutrophil chasing a bacterium. Top right:** Hybrid time series in which 16 FILM-interpolated images were generated between real frames. **Bottom left:** Hybrid time series in which 16 CDFI-interpolated images were generated between real frames. **Bottom right:** Hybrid time series in which 16 LDMVFI -interpolated images were generated between real frames. Brightness normalized across the time series for display purposes.

**Video 9. Representative time-lapse microscopy time series** of pancreas-derived organoids collapsing in response to anticancer drugs. **Top right:** Hybrid time series in which 16 FILM-interpolated images were generated between real frames. **Bottom left:** Hybrid time series in which 16 CDFI-interpolated images were generated between real frames. **Bottom right:** Hybrid time series in which 16 LDMVFI -interpolated images were generated between real frames. Brightness normalized across the time series for display purposes.

**Video 10. Representative time-lapse microscopy time series of an *Ashbya gossypii* nucleus being tracked** consistently in the hybrid time series derived from the FILM and CDFI models but not in the real time series or the hybrid time series derived from the LDMVFI model. Tracking errors in the real time series start at t =26 min or 20 sec in the video.

**Video 11. Representative time-lapse microscopy time series of an *E. coli* bacterium being tracked** consistently longer in the hybrid time series derived from the FILM, CDFI, and LDMVFI models but not in the real time series. Tracking errors in the real time series start at t =30 min or 9 sec in the video.

**Video 12. Representative time-lapse microscopy time series of an *S. cerevisiae* cell being tracked** consistently in the hybrid time series derived from the FILM and CDFI models but not in the real time series, or the hybrid time series derived from the LDMVFI model. Tracking errors in the real time series start at t =276 min or 35 sec in the video, while tracking errors in the LDMVFI time series start at t =348 min or 46 sec in the video.

**Video 13. Representative time-lapse microscopy time series of a Rat Basophilic Leukemia (RBL) cancer cell being tracked** consistently in the hybrid time series derived from the CDFI, FILM, and LDMVFI models, but not in the real time series. Tracking errors in the real time series start at t =200 min or 15 sec in the video.
